# Systematic comparison of experimental assays and analytical pipelines for identification of active enhancers genome-wide

**DOI:** 10.1101/2021.06.02.446833

**Authors:** Li Yao, Jin Liang, Abdullah Ozer, Alden King-Yung Leung, ENCODE Consortium, John T. Lis, Haiyuan Yu

## Abstract

Mounting evidence supports the idea that transcriptional patterns serve as more specific identifiers of active enhancers than histone marks^1,2^; however, the optimal strategy to identify active enhancers both experimentally and computationally has not been determined. In this study, we compared 13 genome-wide RNA sequencing assays in K562 cells and showed that the nuclear run-on followed by cap-selection assay (namely, GRO/PRO-cap) has significant advantages in eRNA detection and active enhancer identification. We also introduced a new analytical tool, Peak Identifier for Nascent-Transcript Sequencing (PINTS), to identify active promoters and enhancers genome-wide and pinpoint the precise location of the 5’ transcription start sites (TSSs) within these regulatory elements. Finally, we compiled a comprehensive enhancer candidate compendium based on the detected eRNA TSSs available in 120 cell and tissue types. To facilitate the exploration and prioritization of these enhancer candidates, we also built a user-friendly web server (https://pints.yulab.org) for the compendium with various additional genomic and epigenomic annotations. With the knowledge of the best available assays and pipelines, this large-scale annotation of candidate enhancers will pave the road for selection and characterization of their functions in a time-, labor-, and cost-effective manner in the future.

---

Regulation of transcription is a synergetic process that requires both trans-regulatory factors, like transcription factors, and cis-regulatory elements, like promoters and enhancers. In contrast to promoters, which initiate transcription in their proximal regions to produce stable RNA products, enhancers regulate transcription of their target gene(s) in a distal manner. Certain epigenomic signatures (enrichment of H3K4me1 and H3K27ac, high chromatin accessibility, and CBP/p300 binding) are considered to be defining features of active enhancer loci^3,4^. However, studies also revealed that enhancers could themselves produce relatively short-lived divergent transcripts, called enhancer RNAs (eRNAs)^5,6^. More recent studies further showed that distal divergent transcription events are more reliable marks for active enhancers than epigenomic signatures^1,2^. Recently we have proposed^7,8^ and later experimentally verified^2^ the basic unit of active enhancers that are defined by the transcription start sites (TSSs) of the divergent eRNA transcription, and delimited by the promoter-proximal Pol II pause sites flanking these TSSs. Therefore, to identify active enhancers genome-wide in any given sample, it is critical to detect eRNAs and their TSSs with high sensitivity and specificity.

eRNAs are usually in extremely low abundance in cells due to their short half-lives. Therefore, conventional RNA-seq experiments capture eRNAs with very low efficiency overall^5^. Recently, two categories of genome-wide RNA sequencing assays have been developed, focusing either on TSSs or on the actively-transcribing polymerase positions (Fig. 1a). We named the 8 assays (GRO^9^/PRO-cap^10^, CoPRO^8^, Start-seq^11^, CAGE^12^, RAMPAGE^13^, NET-CAGE^14^, csRNA-seq^15^, and STRIPE-seq^16^) from the former category as 5’ assays, because these assays enrich for active 5’ TSSs of promoters and enhancers (Fig. 1a). We also named the 5 assays (GRO-seq^17^, PRO-seq^10^, mNET-seq^18^, Bru-seq^19^, and BruUV-seq^20^) from the latter category as 3’ assays, because they are designed to identify 3’ ends of the RNA transcripts (Fig. 1a). To enrich for RNA populations of interests, these assays implement various experimental strategies, including nuclei/chromatin isolation^8–11,18^, nuclear run-on^8–10^ or metabolic labeling^19,20^ with biotin or bromo-tagged nucleotides and affinity purification, Pol II immunoprecipitation^18^, size selection^15,16^, and enzymatic elimination of non-capped RNAs^8–10,16^, or chemical tagging of capped RNAs^12–14^. We summarized key experimental steps of both 5’ and 3’ assays in Fig. 1b. In fact, the list of all assays compared here, plus total RNA-seq^21,22^, have all been utilized in some capacity to identify enhancer elements. However, considering that most of these assays are not specifically designed to capture eRNAs, caution should be taken when exploiting the resulting data for identification of active enhancers or when deciding on the appropriate assay to use. Here, using datasets produced in the same ENCODE Tier 1 cell line, K562, we performed a comprehensive evaluation of the reliability and sensitivity in detecting eRNAs (thus identifying active enhancers genome-wide) across all available RNA sequencing assays, including seven 5’ assays (no K562 Start-seq dataset is available), five 3’ assays, and total RNA-seq (as the outgroup).

**Fig. 1.**
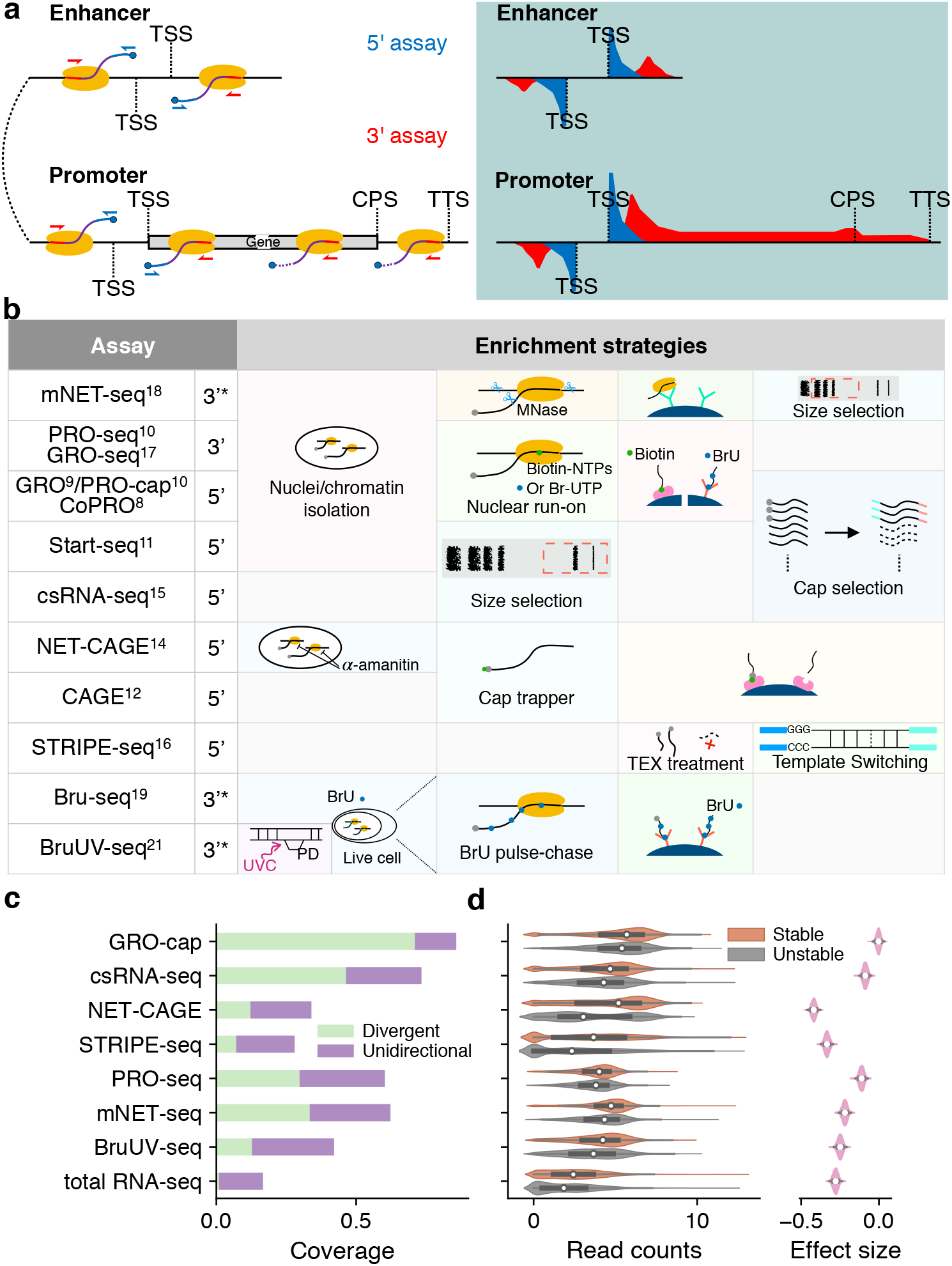
Comparison of currently available assays for detecting eRNAs. **a**, Schematics of enhancer and promoter/gene transcription by RNA Pol II (left panel) and characteristic profiles of 5’ and 3’ assays (right panel, light-blue shaded area). Black lines represent genomic DNA; nascent RNAs are purple curved lines with 5’ and 3’-end colored blue and red, respectively, with light-blue spheres as caps and yellow ovals indicate RNA Pol II. Arrows indicate the direction of sequencing reads, 5’ assay in blue, and 3’ assay in red. Representative read density profiles are colored accordingly in blue and red for 5’ and 3’ assays, respectively. TSS: transcription start site; CPS: cleavage polyadenylation site; TTS: transcription termination site. **b**, Enrichment strategies used by different 5’ and 3’ RNA assays. TEX: terminator exonuclease. * indicates that 3’ RNA ends can only be estimated approximately by these assays. A detailed description is available in Supplementary Notes. **c**, The capability of different assays to capture validated enhancers (true enhancer set). All libraries were downsampled to the same sequencing depth. “Unidirectional” and “divergent” indicate the detection of eRNAs originated from either one or both strands of the enhancer loci, respectively. **d**, Differences in read coverage among stable and unstable transcripts. GRO-cap has the highest coverage of both stable and unstable transcripts, and the least preference toward stable transcripts. Read counts were log(*n* + 1) transformed; preferences (effect sizes) were evaluated as Cohen’s *d*.

Because these assays were initially designed for different purposes, various computational tools were developed for exploring and interpreting the raw experimental data—for example, Tfit^23^, dREG^24,25^, and dREG.HD^26^ were developed to identify transcriptional regulatory elements (TREs) from some 3’ assays, including GRO-seq and PRO-seq; FivePrime^27^ (based on paraclu^28^), GROcapTSSHMM^7^, and HOMER^15^ were introduced for analyzing data from CAGE, GRO-cap, and csRNA-seq, respectively. While all these tools can potentially be used to identify eRNA transcription and active enhancers, there has not been a systematic evaluation and comparison of their performance with datasets generated by the aforementioned experimental assays.

In this study, we systematically examined 13 experimental assays in terms of their sensitivity and specificity for capturing eRNAs. We also developed a novel computational tool, Peak Identifier for Nascent-Transcript Sequencing (PINTS), which is designed to identify enhancer candidates from all of these assays. Moreover, by comparing PINTS with 8 other widely-used computational tools, we found that PINTS gave the highest overall performance pertaining to robustness, sensitivity, and specificity, especially when analyzing data from 5’ RNA sequencing assays. Finally, we constructed a comprehensive enhancer candidate compendium for 120 cells and tissues using the robust and unified definition of active enhancers based on detected eRNA TSSs genome-wide^1,2,7^, and developed an online web server (https://pints.yulab.org/) to navigate, prioritize, and analyze enhancers based on a wide range of genomic and epigenomic annotations. We expect our enhancer compendium will be a valuable resource to the research community for the effective selection of candidate enhancers for further functional characterization in future studies.

## Results

### 5’ assays enriching short and/or capped RNAs demonstrate higher sensitivity in eRNA detection with GRO-cap being the most sensitive assay

To perform a quantitative comparison of eRNA detection sensitivity, we first normalized all libraries by down-sampling them to the same sequencing depth as the library with the lowest depth (18.9 million mappable reads, Supplementary Table 1). We then conducted a pairwise comparison of their coverage at 741 previously identified super-enhancers^29^. The result shows that assays using nuclear run-on followed by cap-selection (GRO-cap^7^/PRO-cap, CoPRO^8^) have a higher coverage over all other assays (Cohen’s *d* of log-transformed coverage: 0.3926~1.1642). Since super-enhancers can span several kilobase pairs in the genome and are believed to be composed of clusters of enhancers, we also compared the assay sensitivity in a higher resolution manner by examining their coverage in 803 (635 intergenic, 113 intronic, and 55 others) bona fide enhancers (referred to as “true enhancer set” in this study, Supplementary Table 2) validated by CRISPR/Cas9-mediated deletion and CRISPRi in K562 cells^30–38^. With the same sequencing depth, GRO-cap ranks first in terms of sensitivity: it covers 85.80% of true enhancers (70.98% divergent: ≥ 5 reads detected from both strands and 14.82% unidirectional: ≥ 5 reads detected only on one strand; Fig. 1c and Supplementary Fig. 1a). csRNA-seq comes in second place with 73.35% (46.33% divergent and 26.40% unidirectional) coverage of these validated true enhancers (Fig. 1c and Supplementary Fig. 1a).

We further evaluated the sensitivity of these assays by their ability to capture unstable transcripts. eRNAs are usually less stable compared to RNAs of other types, as seen in the decay rates^39^ of transcripts originated from 514 enhancers from the true enhancer set, which are significantly more rapid than those of mRNAs (*p*-value: 1.022e-86, Cohen’s *d*: 0.9469). We then used as the cutoff between stable and unstable transcripts the 95^th^ quantile of decay rates of mRNAs and surveyed the distribution of read counts captured in the two categories among all assays. Consistent with our conclusion above, GRO-cap has the smallest differences in read coverage between stable and unstable transcripts [Cohen’s *d*: −0.0028, 95% CI: (−0.0328, 0.0239)], indicating assays utilizing nuclear run-on followed by cap-selection have the greatest ability to enrich unstable transcripts, which is of particular importance for detecting eRNAs (Fig. 1d and Supplementary Fig. 1b). We evaluated the effects of technical artefacts, including strand specificity and internal priming, and our results suggest all libraries have great strand specificity (average: 0.9797, SD: 0.0209) and low internal priming rates (Supplementary Notes).

### Sequencing reads from gene bodies and small RNAs in 3’ assays contribute to lower sensitivity in eRNA detection

For the two families of assays that we compared in this study, we noticed that in general, 5’ assays are more sensitive in detecting eRNAs than 3’ assays, even for assays that use very similar enrichment strategies (Fig. 1b). For instance, while both GRO-cap and PRO-seq employ similar nuclear run-on procedures, there is a 41.22% difference between their divergent coverage of the true enhancer set (Fig. 1c). When inspecting genome-wide distribution of reads (Fig. 2a and Supplementary Fig. 2a), we noticed that 3’ assays have significantly higher proportions of reads coming from gene body regions (mean of 3’ assays: 65.62%, mean of 5’ assays: 13.01%, *p*-value from Mann-Whitney *U* test: 0.0029), which is not surprising as they are designed to reveal all actively transcribing RNA polymerases, whereas 5’ assays are specifically designed for identification of TSSs. Because eRNA transcription is on average much lower than that of genes^7^, such a high portion of gene body reads in 3’ assays dilute the signal from eRNAs and significantly lower their sensitivity in detecting active enhancers. As shown in Fig. 2b, 3’ assays detect a significant number of reads in the FAM89A gene body, whereas 5’ assays only have reads in the promoter regions of the FAM89A gene. As a result, almost all 3’ assays (except PRO-seq) have no discernable signal at a distal enhancer locus near the FAM89A gene that was validated by CRISPRi^35^(Fig. 2b). Another potential problem for 3’ assays is that reads that are mapped to intergenic or intronic regions could be derived from either eRNAs, the unprocessed precursors of RNAs from other categories (e.g., pre-mRNAs), or read-through from an upstream transcription event (Supplementary Fig. 2b), which further reduces their sensitivity for detecting eRNAs.

**Fig. 2.**
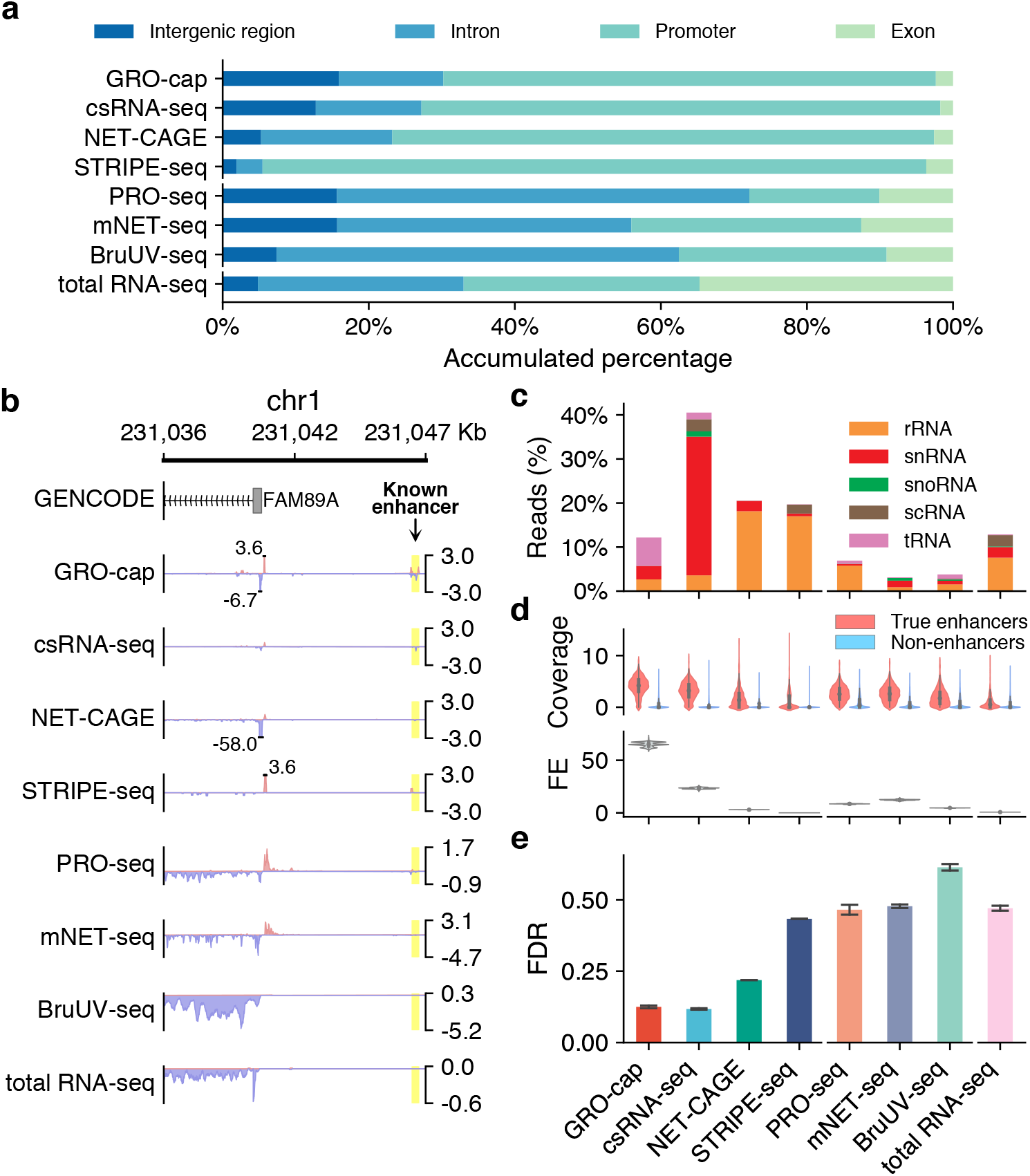
Characterization of factors affecting assay sensitivity and evaluation of assay specificity in eRNA detection. **a**, Genome-wide distribution of sequencing reads originated from intergenic regions, introns, exons, and promoters detected by different assays. **b**, A genome browser snapshot of a gene (FAM89A) and its enhancer (highlighted in yellow), demonstrating the different patterns of signals captured by 5’ (enriched in the promoter and enhancer regions) *vs*. 3’ (enriched in the promoter and gene body regions) assays. Signals are normalized by reads per million (RPM). **c**, Proportion of mappable reads from different assays originated from various abundant RNA families. **d**, Specificity in detecting eRNAs. Top track: differences of read coverage among true enhancer and non-enhancer loci, a pseudo-count (1) was added to each locus, and the coverage was log-transformed. Bottom track: signal-to-noise ratios depicted in forms of fold enrichment (FE), the results are based on bootstrapped samples, median statistics are used to calculate the fold changes. **e**, False discovery rates (FDRs) estimated by the overlap between the top 10,000 genomic bins and the reference (true and non-enhancer) loci. Pseudoreplicates were used (N=3); error bars represent SD.

Most of the RNA transcripts in living cells belong to the families of highly abundant RNAs, e.g., rRNAs and snRNAs. When capturing eRNAs with next-generation sequencing-based methods, a successful exclusion of RNAs of these high-abundance families from the sequencing libraries during the preparation process would greatly enhance the efficiency and sensitivity of eRNA identification. We compiled a comprehensive list of high-abundance RNAs in human cells by incorporating annotations from GENCODE^40^, RefSeq^41^, and RMSK^42^. Based on this list, an average of 7.124% of the mappable reads in each assay originated from rRNAs despite most of the assays employed strategies to reduce rRNA inclusion (Fig. 2c and Supplementary Fig. 2c). By simulating a rRNA-depleted BruUV-seq library, we found that a complete depletion of rRNAs could contribute to a 1.76-fold boost in detecting eRNAs (Supplementary Fig. 2d).

When the assay specifically enriches for short transcripts, like csRNA-seq, a relatively large proportion (31.49%) of the mappable reads were found to have originated from snRNAs (Fig. 2c and Supplementary Fig. 2c). Detection of such a disproportionally large fraction of snRNAs (Fig. 2c) suggests a potential contamination in the sequencing library from the splicing intermediates. To test the possibility, we calculated the signal densities at all the splice sites in the human genome according to GENCODE annotation (v24)^40^. As shown in Supplementary Fig. 2d, csRNA-seq does detect more signals at the splicing junctions compared to all the other assays.

### 5’ assays have better specificity in detecting active enhancers, with GRO-cap having the best performance

While genomic regions with detectable transcriptional events account for 75% of the human genome^43^, many of these events are considered to be spurious transcriptional noise^44,45^ because of their extremely low transcript yields compared to mRNAs and of the intrinsic promiscuity of RNA Pol II under certain circumstances^46^. Therefore, it is critical to detect and differentiate the reads that originated from spurious transcription in these assays. To that end, we collected non-enhancer loci from eight Massively Parallel Reporter Assay (MPRA)^47–51^ and Self-Transcribing Active Regulatory Region Sequencing (STARR-seq)^2,52,53^ studies and further removed elements overlapping with predicted Enhancer-Like Sequence (ELS) or Promoter-Like Sequence (PLS) from candidate cis-regulatory elements (cCRE) annotations^54^ to generate a set of 7,097 loci (referred to as the “non-enhancer set”, Supplementary Fig. 2f, Supplementary Table 3). We observed that signal intensities in the true enhancer set are often higher than that in the non-enhancer set (Fig. 2d and Supplementary Fig. 2g), with GRO-cap having the highest signal-to-noise ratio (64-fold enrichment, Fig. 2d and Supplementary Fig. 2g).

We also calculated false discovery rate (FDR) for each assay based on the overlap of their reads with both the true enhancer and non-enhancer sets. We found that 5’ assays generally have lower FDRs than those of 3’ assays (5’ assay mean: 0.2698 *vs*. 3’ assay mean: 0.5253, *p*-value from Mann-Whitney *U* test: 5.089e-5, Fig. 2e and Supplementary Fig. 2h), with GRO-cap having the lowest FDR (mean: 0.1250, SD: 0.0046).

### A novel computational tool for identifying active enhancers and promoters from 5’ assays: Peak Identifier for Nascent-Transcript Sequencing (PINTS)

High-throughput sequencing-based bioassays rely on sophisticated analyses to identify meaningful information from raw data, so it is essential to choose the appropriate computational tool to do the job. Two categories of tools are currently available for signal processing. Tools in the first category predict entire transcription units, they are primarily used for 3’ assays and often use a set of cutoffs for fold-changes, e.g., HOMER (GRO-seq)^55^ or Hidden Markov Models^19,56^, e.g., groHMM, to determine the start and end positions of transcription units. Tools in the second category usually identify narrower regions for potential regulatory elements (mainly promoters and enhancers, often referred to as peak callers) and include GROcapTSSHMM^7^, dREG^24^, Tfit^23^, dREG.HD^26^, TSScall^11^, HOMER (csRNA-seq)^15^, and FivePrime^27^ (based on Paraclu^28^). Based on previous studies, the peak of divergent TSSs, which is associated with eRNA transcription, is an effective mark for active enhancers^1,2^. Therefore, in this comparative study, we focused on the second category of computational tools.

To achieve a higher resolution in identification of transcriptional regulatory elements, a common practice is to only look at the ends of the transcripts captured by these assays. When such a practice is adopted, two critical issues emerge when evaluating the statistical significance of peaks (i.e., TSSs), especially for the 5’ assays. First, when only taking the transcript ends into account, the fraction of zeros (no mapped reads per basepair) in local background increases, which deflates read density in the local background and thus inflates the statistical significance of candidate TSSs and resulting in false positives. Second, multiple TSSs can localize in close proximity in the genome and therefore inflate the estimation of read density in the local background, resulting in diminished statistical significance for all TSSs in that locus and leading to false negatives in TSS detection. To address these issues, we developed Peak Identifier for Nascent-Transcript Sequencing (PINTS), which uses zero-inflated Poisson models to evaluate local read densities and employs interquartile range (IQR)-based refinement to ameliorate false negatives by conditionally masking candidate TSSs in the local background (Fig. 3). PINTS was inspired by MACS2^57^ with modifications specifically implemented for identification of eRNA TSSs from genome-wide RNA sequencing assays (especially 5’ assays). After evaluating the significance of each TSS, PINTS defines TREs as divergent TSS pairs that are within 300bp from each other, as suggested by previous studies^2^ (Supplementary Fig. 3a, Methods). We identify candidate enhancers as the distal TREs that are farther than 500 bps away from known protein-coding gene TSSs^2^.

**Fig. 3.**
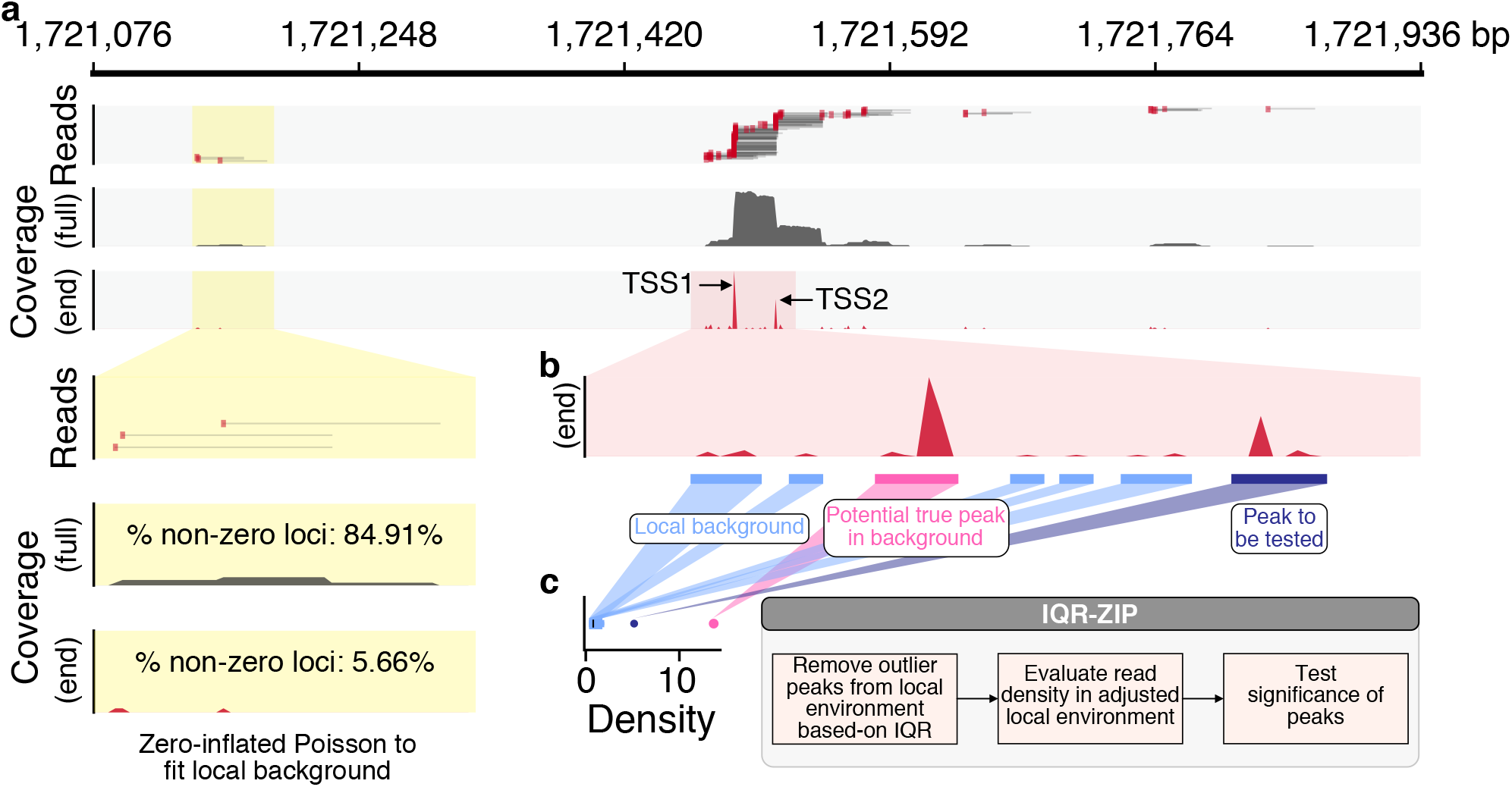
Peak Identifier for Nascent-Transcript Sequencing (PINTS): a novel computational tool for identifying active enhancers and promoters from 5’ assays. **a**, Improvement of TSS identification resolution by focusing only on read ends and using zero-inflated Poisson (ZIP) models to fit local background to address the significantly increased sparsity of signals. The thin grey lines indicate sequencing reads with the 5’ ends highlighted in red. **b**, The existence of other potential true peaks (pink) elevates the estimation of read density in the local background. **c**, Schematic plot shows how IQR-ZIP works. The blue box shows the read density distribution of the local background; the purple dot shows the density of the peak to be tested; the pink dot shows the density of a potential true peak close to the peak to be tested, whose read density is a clear outlier and thus excluded from local background estimation.

### Most peak callers can identify active enhancers from high-throughput datasets, but their resolution and computational requirements vary significantly

Candidate enhancer loci identified by peak caller algorithms should share the same features as true enhancers with characteristic epigenomic marks (DHS, H3K27ac, H3K4me3, and H3K4me1) and transcription factor (CBP/p300, GATA1, and CTCF)-binding sites. Indeed, we found that candidate enhancers identified by most tools recapitulate these features of true enhancers (Fig. 4a, b and Supplementary Fig. 3a for GRO-cap and Supplementary Fig. 3b~1 for all other assays). However, the distribution pattern of epigenomic marks and TF-binding sites of candidate enhancers identified by MACS2^57^, a widely used peak caller for analyzing ChIP-seq data, is remarkably distinct from those of the true enhancers, suggesting the default shifting model of MACS2 optimized for ChIP-seq data may not be suitable for identifying eRNA TSSs of active enhancers.

**Fig. 4.**
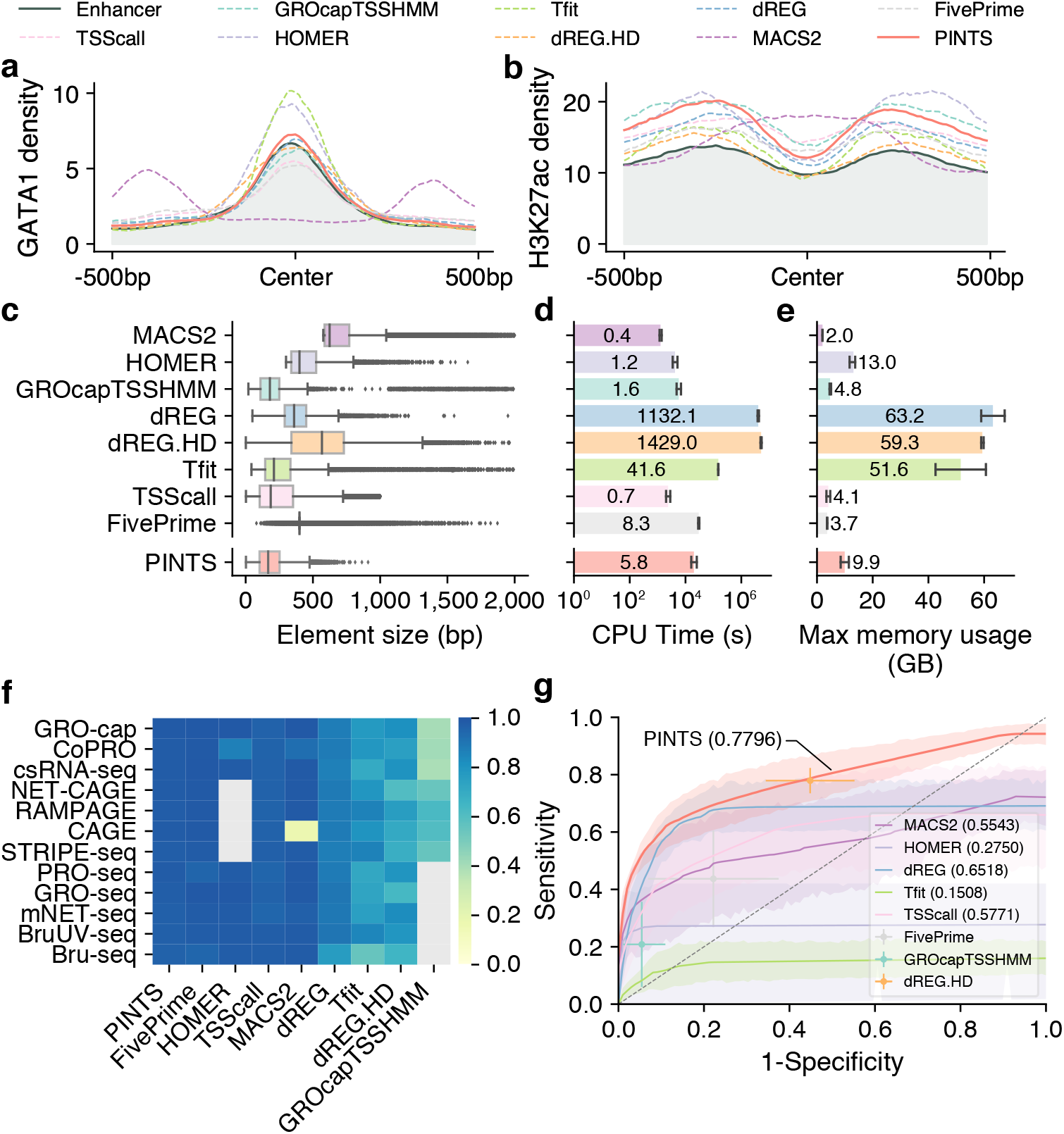
PINTS achieves the best balance among resolution, robustness, sensitivity, specificity, and computational resources required. **a** and **b**, Profiles of GATA1 binding sites and H3K27ac pattern in true enhancer regions and distal TREs identified by different peak callers. **c**, Distribution of element sizes identified by different tools. **d**, CPU time consumed by peak callers to identify elements from various 5’ assay libraries. Average CPU time labeled inside each bar is in the unit of hours (N=6). The x-axis is in log(n + 1) scale. **e**, Maximum memory usage during peak calling from 5’ assay libraries. Average maximum usage labeled inside or on top of each bar is in the unit of gigabytes (N=6). For **d** and **e**, error bars represent SD. **f**, Robustness of peak calls made by different tools. Libraries were mapped to both hg19 and hg38, and robustness was measured as the Jaccard index between calls from hg19 and hg38 (lifted over). Cells colored in gray indicates either that the tool cannot be applied to the corresponding assays or that one or more required datasets are not available. **g**, Aggregated ROC curves for each peak caller on all 5’ assay datasets. The solid lines represent the mean values; the corresponding shaded areas show the 95% confidence interval of the means (via bootstrap). For tools where ROCs cannot be calculated, solid dots represent their performance with default parameters, error bars show SD (see Methods).

We further surveyed the size distribution of these elements as an indication of peak-calling resolutions. We found that PINTS, GROcapTSSHMM, Tfit, and TSScall achieve higher resolution (average peak size of 185-258 bp) than other tools when analyzing 5’ assay data (Fig. 4c and Supplementary Fig. 3m for all other assays). Notably peaks called by dREG.HD and MACS2 range between 548-751 bp in size, whereas dREG, FivePrime, and HOMER peaks range between 381-460 bp.

The amount of computational resources required by a peak caller greatly affects its general applicability. Here, we compared the total amount of CPU time and peak memory usage that each peak caller requires for identifying divergent elements from 5’ assays (Fig. 4d and 4e). Based on our calculation, it is feasible to run PINTS, MACS2, HOMER, GROcapTSSHMM, TSScall, and FivePrime on a typical personal computer.

### PINTS achieves high overall robustness, sensitivity and specificity in detecting TREs from 5’ assays

A common strategy in functional studies of enhancers is to compare active enhancers across different conditions and diseases^21,22^, which requires the *in silico* enhancer-predicting tools to be robust against biological and experimental variances. To test the upper bound of robustness for the aforementioned tools, we performed peaks calls using these tools on datasets from 12 assays by aligning each dataset to two commonly-used builds of human reference genome: hg19 and hg38. Because there is no variation in the RNA sequencing datasets themselves and the two reference genome builds are very similar (hg38 has 0.09% more ungapped non-centromeric sequences than hg19, only 0.17% of ungapped hg19 sequences are not in hg38^58^, Supplementary Fig. 4a), we expect that the differences in peak calls using these two different genome builds should be minimal across all datasets. Surprisingly, we found that by simply changing the reference genome builds, half of them have Jaccard index smaller than 0.9 for at least one assay (Fig. 4f). Furthermore, we noticed that Jaccard indices were even lower when we tried to evaluate robustness across real technical and biological replicates (average: 0.5068, SD: 0.2462), especially for TSScall and FivePrime, where their robustness is only 0.4013 (SD: 0.1038) and 0.3836 (SD: 0.2028), respectively (Supplementary Fig. 4b). PINTS consistently have great robustness in both cases (average: 0.9761, SD: 0.0081 between hg19 and hg38 genome builds, and average: 0.7279, SD: 0.0515 across replicates).

To evaluate sensitivity and specificity, two other key metrics of performance, we merged the true enhancer set with the promoter regions from GENCODE v24^40^ as the positive set, and non-enhancer loci as the negative set (Method). As shown in Supplementary Fig. 4c, the negative set has distinct patterns of TF-binding motifs and chromatin marks compared to either the true enhancer or the promoter set. We then evaluated each tool’s performances for all 5’ assay datasets (Fig. 4g). The results show that PINTS achieves the best balance between sensitivity and specificity (PINTS mean AUC: 0.7796, SD: 0.0821; mean AUC for the second-best tool dREG: 0.6518, SD: 0.1088). For all of these computational tools, we summarize their key requirements, main characteristics, and applicability to different RNA sequencing assays in Supplementary Fig. 5.

### An enhancer compendium for human cell and tissue types

Having comprehensively evaluated the performance of all available whole-genome RNA sequencing assays and computational tools in identification of candidate enhancers genomewide, we reasoned that available 5’ assay datasets across different human cells and tissues could be analyzed by PINTS to construct a human enhancer candidate compendium. Previous studies have shown that, compared with histone marks, detecting enhancers by divergent eRNA TSSs has advantages in both resolution and specificity^1,2^. Such an eRNA-centric enhancer compendium, in addition to all the available enhancer datasets based on histone marks^54,59^, will be an invaluable resource to better understand gene regulation, to functionally annotate the noncoding genome, and to help prioritize non-coding variants across disease cohorts by their potential impact on enhancer activities^60^. Toward this goal, we applied PINTS to identify candidate enhancers using 5’ assay datasets (i.e., GRO/PRO-cap, Co-PRO, csRNA-seq, NET-CAGE, RAMPAGE, CAGE, and STRIPE-seq) across 33 cell lines, 7 *in vitro* differentiated cells, 35 primary cells, and 45 tissue samples, including all available 5’ assay datasets through the ENCODE portal (Fig. 5a). Such a comprehensive catalog of enhancers across a wide range of human cells and tissues analyzed by the same exact computational pipeline provides an excellent resource to perform meaningful comparative genomic analyses to study the dynamics of enhancers and gene regulation in general, which will help focus on true biological differences while minimizing technical variations. In addition, for 7 human cell lines (K562, GM12878, HepG2, HeLa-S3, MCF-7, H9, and HCT116), we applied all other available tools (FivePrime, HOMER, TSScall, dREG, dREG.HD, and Tfit) to identify candidate enhancers. We believe this unique resource of enhancers in 7 cell lines with multiple 5’ assay datasets analyzed by all available computational tools will greatly help further the studies of enhancers and their key architecture characteristics. All of these candidate enhancer calls are made publicly available through our web server (https://pints.yulab.org) described in detail below. We will regularly update our enhancer compendium as new datasets, especially those in new cell lines or samples, and assays, become available.

**Fig. 5.**
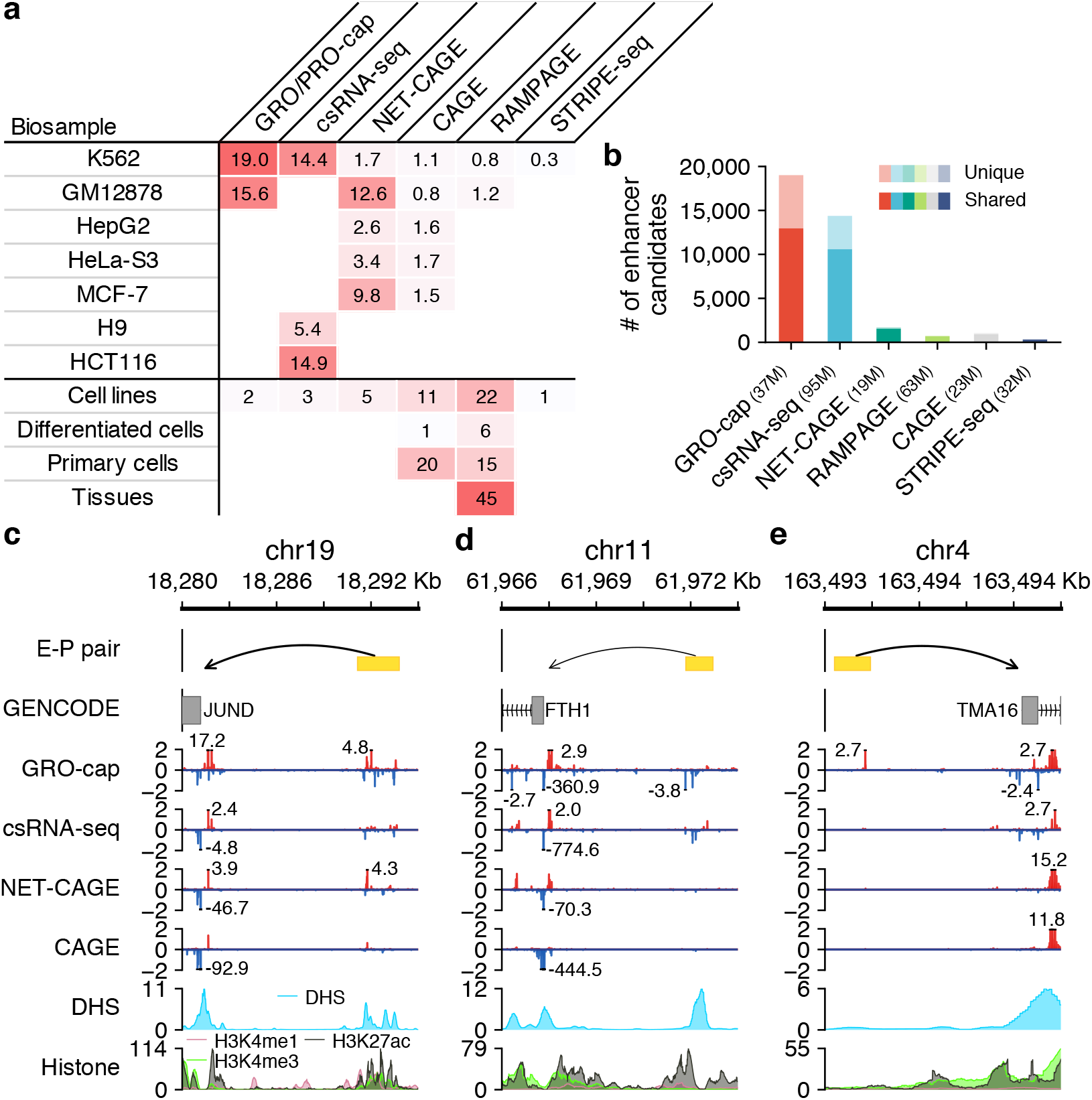
A comprehensive human enhancer compendium. **a**, Summaries of all distal elements identified from different assays with PINTS in 7 cell lines (top) and 131 datasets generated by different assays across 120 biosamples included in our enhancer compendium (bottom). Differentiated cells stand for *in vitro* differentiated cells. **b**, The number of distal elements identified by PINTS from different assays in K562. The dark colors indicate the proportion of shared elements identified from at least one other assay; light colors indicate elements unique to the corresponding assay. The number of mappable reads for each assay is listed in the parentheses. **c**, A known enhancer locus where most of these assays captured signals on both strands. **d**, A known enhancer locus where only GRO-cap and csRNA-seq captured detectable signals. **e**, A known enhancer locus where only GRO-cap captures clear signals. Signal tracks in **c**, **d**, and **e** were normalized by their sequencing depths (RPM).

In human K562 cells where datasets are available from all 5’ assays, our results show that GRO-cap has by far the most number of distal TRE elements (19,006 identified by PINTS with 9,531 unique enhancer calls – not identified by any other assay; the second-best dataset, csRNA-seq, only has 14,375 enhancer calls with 5,048 unique (Fig. 5b). This is not surprising given that GRO-cap has the best sensitivity in detecting eRNA transcription (Fig. 1c and 1d) and the GRO-cap dataset has the second highest read depth (Fig. 5b). We selected three CRISPRi-validated enhancer-promoter pairs^35^ to visualize these differences and showed the variety in signal abundances between all assay datasets (Fig. 5c-e). For example, the enhancer which regulates the JUND gene (Fig. 5c) has decent accessibility and is supported by epigenomic marks, including H3K27ac and H3K4me1. As expected, all four 5’ assays can identify this enhancer. The expression levels of enhancers are not necessarily to be proportional to the levels of epigenomic marks, and for eRNAs whose expression levels are lower (e.g., the enhancer that regulates FTH1 in Fig. 5d), assays that are more effective in capturing unstable transcripts are more likely to recover them. Finally, for the enhancer regulating TMA16, signals from histone marks are quite minimal, but GRO-cap still captures clear signals of eRNA transcription at this locus and readily enables identification of this enhancer (Fig. 5e).

### An online web server for exploring and analyzing enhancers and various genomic and epigenomic features genome-wide

To make it easier for biologists to explore the tens of thousands of candidate enhancers in our compendium and to prioritize these enhancers for further studies, we constructed an online web server (https://pints.yulab.org) that is publicly available, where users can query any human genomic region of their interest in a given biological sample to get a comprehensive list of candidate enhancers detected by any available 5’ RNA sequence assays (i.e., GRO/PRO-cap, csRNA-seq, NET-CAGE, CAGE, RAMPAGE, and STRIPE-seq) in that sample using any of the 7 peak callers (PINTS, FivePrime, dREG, dREG.HD, HOMER, Tfit, and GROcapTSSHMM). We also included detailed annotations for each candidate enhancer, including epigenetic features^54^, core promoter elements^61^, potential transcription factor binding sites^62^, and presence of population variants and ClinVar mutations^63^, as shown in Fig. 6. Users can prioritize enhancers by requiring specific supporting 5’ assay data available, the presence of certain epigenomic features, core promoter elements and transcription factor binding sites, or containing different categories of population variants (common vs. rare) and ClinVar mutations (benign vs. pathogenic). For the transcription factor binding site analysis, our web server will automatically integrate the information from available RNA-seq data to only include those factors that are expressed in the selected cell line.

**Fig. 6.**
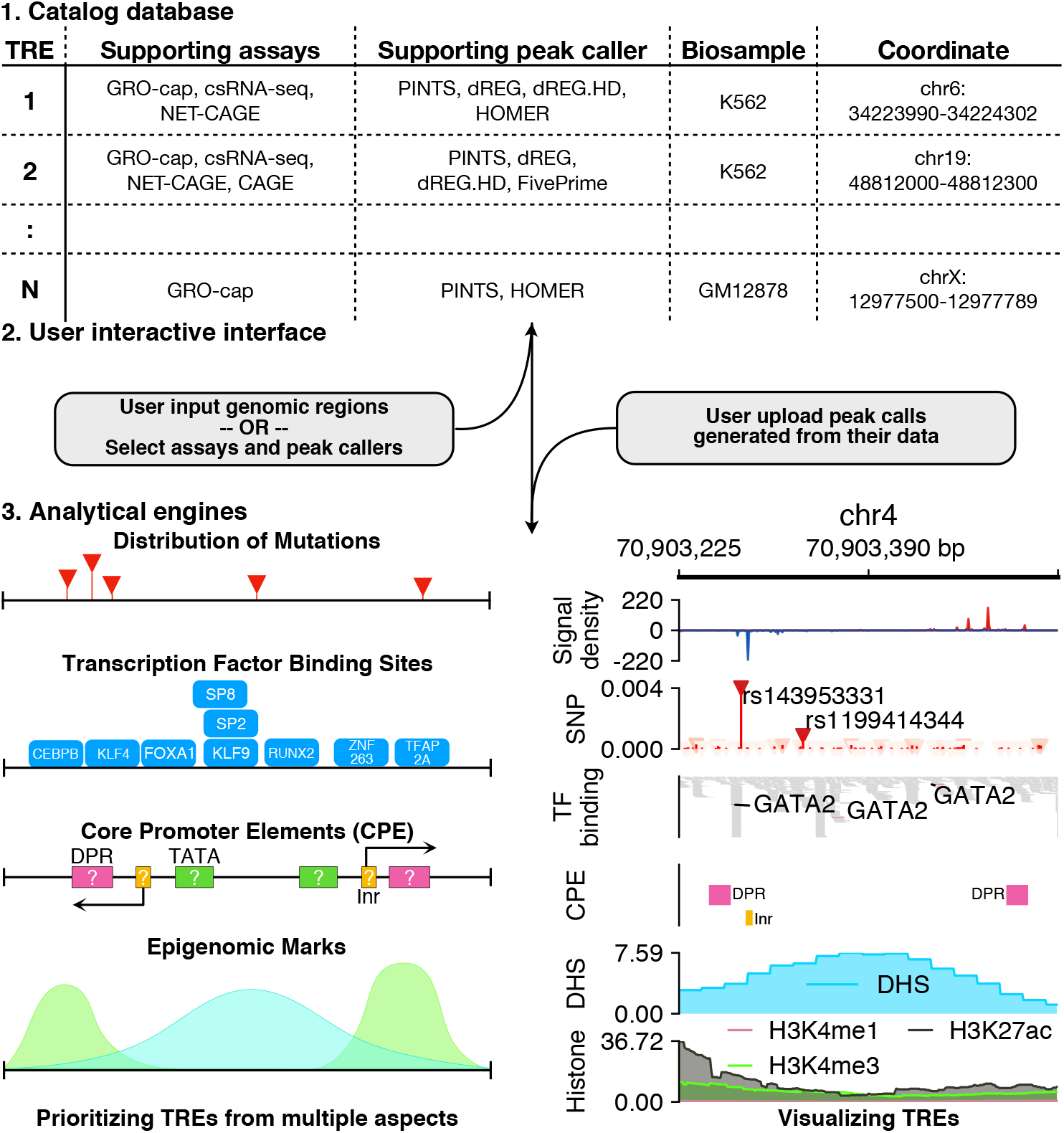
Interactive PINTS web server. The server is composed of three modules: a catalog database containing our human enhancer compendium across 120 biosamples, a user interactive interface, and analytical engines. When the user provides genomic loci of their interest as search queries, previously annotated information related to the loci will be retrieved from the catalog database. The user can further refine the search results by a series of filters, including the supporting assays, callers, the presence of mutations, transcription factor binding sites, core promoter elements, and epigenomic marks. On the other hand, if the user chooses to upload peak calls generated from their own data, the uploads will be analyzed by the analytical engines to provide the same types of annotations as those in the catalog database.

Furthermore, users can upload their own enhancers calls in a human cell line or type, our web server will automatically annotate all of their enhancers calls with the same pipeline as described above. We have integrated epigenomic features and RNA-seq data for 197 human-derived biosamples from the ENCODE project in our web server. For user-uploaded enhancer data in these samples, we automatically refine our annotations by only reporting binding sites of expressed transcription factors, and associating each enhancer with epigenomic features specific to the corresponding sample.

Users can explore all annotations of their selected enhancers via our integrated genome browser; alternatively, they can easily export all annotations to a local machine in plain text format, which greatly facilitates any user-designed downstream analyses.

## Discussion

eRNAs are increasingly being recognized as a critical marker for active enhancers genome-wide^1,2^; however, the optimal strategy (both experimental assays and their analytical pipelines) to detect eRNAs and thus identify enhancer loci has not been unveiled. In this study, we systematically compared 13 *in vivo* genome-wide RNA sequencing assays in K562 cells and showed that 5’ assays are in general more sensitive than 3’ assays to detect eRNAs, because signals will not be diluted by active transcription in gene bodies. One additional, and critical advantage of the 5’ assays is that they reveal the precise location of eRNA TSSs, allowing for high-resolution detection and dissection of enhancer loci genome-wide as demonstrated in our recent work^2^. Overall, our results show that GRO/PRO-cap has the best overall performance in detecting active enhancers in terms of both sensitivity and specificity.

We noticed that when using current computational tools to identify TREs from various RNA sequencing datasets, very minor changes in sample processing could lead to more than 20% of changes in the final results, which brings the robustness of the peak calls into question. To address this issue, we introduced a new tool, PINTS, which was inspired by MACS^57^ but with unique implementations (e.g., using zero inflated Poisson distribution to account for sparsity of only mapping 5’ of all reads) that were designed specifically for defining TSSs precisely when analyzing 5’ RNA sequencing data. Our benchmarks indicate that PINTS achieves the best balance among robustness, applicability, sensitivity, and specificity, especially for 5’ RNA sequencing assays capable of detecting the precise location of eRNA TSSs.

In this study, we used CRISPR/Cas9 and CRISPRi validated enhancers^30–38^ as the positive reference set, and MPRA/STARR-seq negative segments^2,47–53^ as non-enhancers. Although these two sets show quite different epigenomic profiles (Supplementary Fig. 2f), which indicates that our non-enhancer set is depleted of any true enhancers, there might still be a few false negatives in the non-enhancer set because not every enhancer works with every promoter, but in the published MPRA/STARR-seq datasets, only a very small number of promoters were used to test all candidate elements; furthermore, some tested elements might be truncated due to synthesis limitations (<200 bp) or random fragmentation of the genome. However, such cases are not expected to affect our relative ranking of different assays and thus will have minimal impact on our conclusions.

In summary, the comprehensive comparison and thorough analyses of 13 genome-wide RNA sequencing assays and 9 computational tools presented here help better understand the strengths and weaknesses for each assay and tool, with regard to detecting active enhancers *in vivo*. Given the fundamental importance of enhancers in gene regulation and the immense interest in accurately identifying active enhancers in a wide variety of samples across species, our results provide basic guidelines in selecting the proper experimental assays and the correct computational tools for future studies.

Furthermore, we provide a detailed, comprehensive human enhancer compendium for 33 cell lines, 7 *in vitro* differentiated cells, 35 primary cells, and 45 tissues (120 in total). We used a unified definition^2,7^ of enhancers based on the detected divergent pairs of eRNA TSSs (i.e., peak calls from various genome-wide RNA sequencing assays, especially 5’ assays). Such a robust, unified and comprehensive catalogue of enhancers across 120 cell types and tissues is expected to shine light on the mechanism of gene regulation and architectural details of enhancers in general. Precise definition of enhancer element boundaries afforded by 5’ assays like PRO/GRO-cap would alleviate potential concerns regarding whether full-length enhancer elements were selected and tested in follow-up functional studies, and thus improve coverage of elements by eliminating incomplete or ill-defined candidates. Such a well-defined catalogue of enhancers also provides an invaluable resource for follow-up studies to better understand the similarities and key differences in gene regulation across various tissues and conditions, and to identify key enhancers whose malfunctions can lead to specific disorders.

## Methods

### Data preprocessing

All datasets were managed and analyzed with BioQueue^64^ (Supplementary Table 1). Raw reads were preprocessed with fastp^65^ according to the experimental designs reported in original papers. Only reads longer than 14bp were kept for downstream analyses. All processed reads from RNA assays were aligned using STAR^66^ to primary assemblies of human reference genome hg38 (GCF_000001305.15) together with rDNA (U13369.1) with parameters as --outSAMattributes All --outSAMmultNmax 1 --outFilterMultimapNmax 50. For studies using Drosophila cells (dm6) or other specific samples as spike-ins, Drosophila reference genome (dm6) or the corresponding reference sequences used in the original studies were incorporated into the index. To measure the robustness of peak prediction, we also mapped reads to primary assemblies of human reference genome hg19 (from UCSC, sequences from alternative loci/haplotypes were removed the same way as for hg38).

### Determining read coverage among reference regions

Sequencing reads of replicates for the same assay were merged together and downsampled to the same sequencing depth (the same number of mappable reads) for three times using picard with parameter STRATEGY = Chained. These downsampled data were then converted to bed files to calculate the fraction of overlap between the sequencing reads and the reference regions in a strand-specific manner.

### Classifying the transcription units as stable and unstable units with TT-seq

Transcript annotations derived from TT-seq (GSE75792^39^) were downloaded from the GEO database. Transcription units with NA values were discarded. The 95^th^ quantile of estimated decay rates for mRNAs was used as the cutoff between the unstable (above the cutoff) and stable (below the cutoff) transcription units.

### Characterizing the genome-wide distribution of reads

The entire genome was classified into four categories based on the annotations in GENCODE (ver 24)^40^: exonic and intronic regions were defined as in GENCODE, except that any region with overlapping intronic and exonic annotation was considered as exonic; the 500bp regions flanking annotated transcription start sites of protein-coding transcripts were annotated as promoters; all other regions were considered as intergenic. Sequencing reads of various assays were assigned to the categories of promoters, introns, exons, or intergenic regions (in the exact order) if they were aligned to the corresponding annotated regions in the genome.

### Identifying sequencing reads from splicing intermediates

The exact or approximate positions of transcript termini were inferred from the read ends and the abundance of their corresponding transcripts was normalized as RPM for this analysis. A list of annotated splice junctions and their 200bp flanking regions in the human genome was compiled based on GENCODE v24^40^. For each assay, we iterated through this list and recorded normalized read counts at each position. In Supplementary Fig. 2e, both the average of signals and the 95% confidence interval (estimated by bootstrap) of the averages were reported.

### Compiling the true enhancer and non-enhancer sets

The experimentally quantified enhancer activity of various DNA elements were collected from previous studies (enhancers: 938 from CRIPSR^32^ or CRIPSRi^30,31,33–38^; non-enhancers: 20,941 from STARR-seq^2,52,53^ and 17,462 from MPRA^47–51^). Overlapping elements within the same category were merged until the resulting elements overlap with elements in the other category. Non-enhancer loci were excluded in the final set if they 1) were shorter than 250bp; 2) overlapped with PLS or ELS predicted by cCRE^54^, or 3) overlapped with possible promoters (1kb regions flanking TSSs in GENCODE).

### Calculating FDRs for assays

multiBamSummary was used to generate tables of read counts across the genome in 500-bp bins. For each assay, the bins were ranked by counts in them and the top n bins were considered as true signals from the assay (four cutoffs were tested: 5,000, 10,000, 20,000, and 100,000, Supplementary Fig. 2h). If a bin overlapped with a locus in the true enhancer set, then the bin was considered as a true positive (TP); if a bin overlapped with a locus in the non-enhancer set, it was considered as a false positive (FP). The false discovery rates were calculated as:

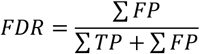

### PINTS

Briefly, read ends are separated based on their mapping directions on the reference genome (forward or reverse), and the read counts are binned into 100-bp windows. Adjacent windows with reads available are merged to avoid splitting potential TRE elements. Within each window, the algorithm first finds peak seeds using a prominence-based approach. Then with a maximum-scoring pairing strategy^28^, the nearby seeds will be merged together as peak candidates if the density after merging meets the following condition:

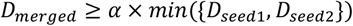

The default value for *α* is 1, and PINTS’s resolution can be further fine-tuned by incorporating reference annotations. For example, when the transcript annotation is available, PINTS will try to avoid the overlap of peak candidates with more than one transcript.

Next, to address the significantly increased sparsity of signals when only the read ends are taken into account, the Expectation-Maximization algorithm is used to fit zero-inflated Poisson (ZIP) models to both peak candidates and their neighborhood regions (λ for read density, π for the proportion of zeros that are not coming from a Poisson process), the probability mass function of these models has the following form:

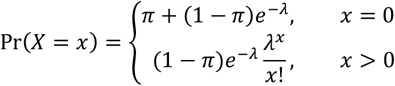

Assume an unobservable latent random variable *z_i_*, for a window *X* of *I* observations, the complete log-likelihood is proportional to:

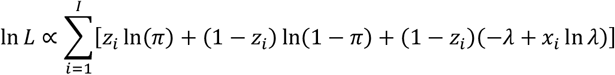

In E-step at the (r + 1)th iteration, *z_i_* is estimated by its conditional expectation:

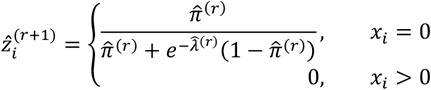

In M-step, given 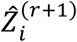 the estimations of π and λ are updated as follows:

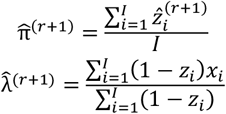

An interquartile range (IQR)-based refinement is applied before fitting ZIP models to the neighborhood regions. In this case, if certain peak candidates s in a local environment are considered as outliers by IQR (their densities are above Q3 + 1.5 × *I*QR, where IQR = Q3 – Q1), these candidates will be masked. For libraries with low sequencing depth, instead of simultaneously masking all outlier peak candidates in the local background, PINTS masks one peak candidate at a time and calculates the resulting peak density in the local background. The process will be reiterated until PINTS either identifies the outlier candidates or reports the nonexistence of such outliers. The estimated densities are then used to determine the statistically significant peaks, which are further categorized into divergent peak pairs (peaks on opposite strands and within 300 bp) and unidirectional peaks.

### Generating peak calls with existing tools

Peak calls for different assays were made using default parameters for other existing peak callers with the following exceptions. For MACS2, --keep-dup all was set so that reads mapped to the same loci would be kept. For FivePrime, parameters *D_min_*, *P_min_*, and *S_min_* were optimized according to the sequencing depth of corresponding libraries, and both divergent TSS calls and enhancer calls were combined as the final output. All tools were allowed to create up to 16 threads/subprocesses if they allow multithreading or parallel computing. For peak callers that do not primarily identify divergent peaks, unidirectional peaks were paired as long as they were within 300 bp and on opposite strands. Maximum memory usage and CPU time (sum of all threads) were monitored with the help from BioQueue^64^. All peak calls were generated on machines with Intel Xeon Gold 6152 CPU @ 2.10 GHz with 88 cores, 1006 GB of RAM running CentOS 7.6.1810.

### Evaluating the systematic biases of different peak calling methods

For each assay, divergent elements were identified using all applicable peak callers, including PINTS. To accommodate the size difference in these elements as well as elements in the true enhancer set, a 1,000-bp region centered around the midpoint of each element was used to evaluate the performance of different methods.

### Evaluating the upper bound of peak caller robustness

Sequencing reads were aligned to another popular reference genome sequence, hg19, and divergent elements were identified accordingly with different peak callers. Peak calls generated from both genome releases were cross lifted using UCSC’s liftover and the average between the two Jaccard indices was considered as the upper bound robustness (UBR):

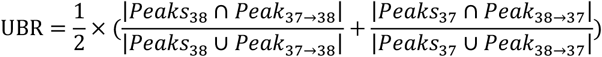

### ROC

For each assay, any element from the true and non-enhancer set was filtered out if there were no sequencing reads aligned to both strands of the element. The positive set was composed of an equal number of randomly sampled promoters (1kb regions flanking TSSs in GENCODE v24) from expressed genes and the filtered true enhancers. The negative set was composed as the filtered non-enhancers. ROCs were generated by calculating the number of divergent elements overlapping with the positive and negative sets under different cutoffs of scores: *p*-values of peaks for PINTS and MACS2, FDRs for TSScall, output SVR scores for dREG, likelihood ratio scores for Tfit, and peak scores for HOMER (findcsRNATSS.pl and findPeak-style TSS). For dREG.HD, GROcapTSSHMM, FivePrime, and HOMER (GRO-seq), since there are either no scores or multiple scores returned in the final output, the sensitivity and specificity was evaluated and reported with their default parameters.

### PINTS web server

The PINTS web server is powered by the Django web framework. dbSNP v153^67^ was used for annotating mutations among TREs; JASPAR 2020^62^ was employed for annotating transcription factor registries and only TFs expressed in the corresponding the biosample (based on RNA-seq data from ENCODE) were included (see Supplementary Table 4 for list of datasets used and accession information). cCRE v2^54^ was used for epigenomic annotation. Core promoter elements were annotated using the following strategy: for each major TSS (+1), the portal annotated the elements as having an initiator or an initiator-like element when the sequence of −3~+3 matches BBCABW; or the sequence of+1~+2 matches YR, respectively. The TATA box (−32~-21) and DPR elements (+17~+35) were identified using the previously published SVR model^61^.

## Acknowledgments

Computation was performed on a cluster administered by the Biotechnology Resource Center at Cornell University. We thank members of the Yu and Lis laboratories, and the ENCODE Consortium (specifically Ali Mortazavi, Mats Ljungman, and Jill E. Moore) for helpful discussions and guidance; Hongya Zhu for her suggestions on concept visualization. This work was supported by a grant from the National Institutes of Health (HG009393 to J.T.L. and H.Y.). L. Y. was supported by the Cornell Presidential Life Sciences Fellowship.

## Author contributions

Conceptualization: L.Y., J.T.L., and H.Y.; Methodology: L.Y.; Software: L.Y.; Formal analysis: L.Y.; Investigation: J.L.; Data curation: L.Y., J.L., and A.K.Y.L.; Writing – Original Draft: L.Y., and J. L.; Writing – Review & Editing: J.L., A.O., J.T.L., and H.Y.; Visualization: L.Y., J.L., A.O., and H.Y.; Supervision: J.T.L., and H.Y.

## Data and materials availability

The source code of PINTS is publicly available at https://github.com/hyulab/PINTS, scripts used to generate results that are reported in this study can be retrieved from https://github.com/hyulab/PINTS_analysis. All sequencing data was retrieved from public databases (NCBI GEO and ENCODE portal), lists of accessions are available in Supplementary Table 1 and 2.

## Supplementary Materials

Figures S1-S5

Supplementary Notes

Tables S1-S4

References (S1-S4)

